# Stringent and complex sequence constraints of an IGHV1-69 broadly neutralizing antibody to influenza HA stem

**DOI:** 10.1101/2023.07.06.547908

**Authors:** Qi Wen Teo, Yiquan Wang, Huibin Lv, Timothy J.C. Tan, Ruipeng Lei, Kevin J. Mao, Nicholas C. Wu

**Affiliations:** Department of Biochemistry, University of Illinois Urbana-Champaign, Urbana, IL 61801, USA; Carl R. Woese Institute for Genomic Biology, University of Illinois Urbana-Champaign, Urbana, IL 61801, USA; Center for Biophysics and Quantitative Biology, University of Illinois Urbana-Champaign, Urbana, IL 61801, USA; Carle Illinois College of Medicine, University of Illinois Urbana-Champaign, Urbana, IL 61801, USA

**Author notes:** To whom correspondence may be addressed. (N.C.W.). These authors contributed equally to this work (Qi Wen Teo, Yiquan Wang, Huibin Lv).

## Abstract

IGHV1-69 is frequently utilized by broadly neutralizing influenza antibodies to the hemagglutinin (HA) stem. These IGHV1-69 HA stem antibodies have diverse complementarity-determining region (CDR) H3 sequences. Besides, their light chains have minimal to no contact with the epitope. Consequently, sequence determinants that confer IGHV1-69 antibodies with HA stem specificity remain largely elusive. Using high-throughput experiments, this study revealed the importance of light chain sequence for the IGHV1-69 HA stem antibody CR9114, which is the broadest influenza antibody known to date. Moreover, we demonstrated that the CDR H3 sequences from many other IGHV1-69 antibodies, including those to HA stem, were incompatible with CR9114. Along with mutagenesis and structural analysis, our results indicate that light chain and CDR H3 sequences coordinately determine the HA stem specificity of IGHV1-69 antibodies. Overall, this work provides molecular insights into broadly neutralizing antibody responses to influenza virus, which have important implications for universal influenza vaccine development.

## INTRODUCTION

Influenza viruses pose a constant threat to public health, resulting in substantial morbidity and mortality with approximately 500,000 deaths worldwide each year.^1^ Although vaccination remains the foremost measure for preventing and controlling influenza virus infection, the effectiveness of the annual seasonal influenza vaccine has varied widely, ranging from 19% to 60% over the past decade.^2^ This variability is largely due to the continuous antigenic drift of human influenza virus, especially at the immunodominant head domain of hemagglutinin (HA), which can in turn lead to vaccine mismatch in some influenza seasons.^3, 4^ In addition, the current seasonal influenza vaccine is designed to protect against influenza A H1N1 and H3N2 subtypes, as well as influenza B virus, but not avian influenza A subtypes with zoonotic potential, such as H5N1 and H7N9. Therefore, significant efforts have been made to develop a universal influenza vaccine that targets the highly conserved HA stem domain,^5, 6^ with an ultimate goal to elicit broadly neutralizing HA stem antibodies like CR9114, which can bind to both group 1 (e.g. H1 and H5) and group 2 (e.g. H3 and H7) HAs as well as influenza B HAs.^7^

Many HA stem antibodies, including CR9114, are encoded by IGHV1-69.^8–10^ Structural analyses have shown that the paratopes of these IGHV1-69 antibodies are dominated by the heavy chain with minimal to no contribution from the light chain.^7, 11–13^ Consistently, diverse light chain germline genes are found among IGHV1-69 HA stem antibodies (e.g. IGKV1-44,^7^ IGLV1-51,^9, 12^ IGLV10-54,^11^ and IGKV3-20^13^). Similarly, the complementarity-determining region (CDR) H3 sequences, which are formed by VDJ recombination, are highly diverse among IGHV1-69 HA stem antibodies.^14^ Although most IGHV1-69 HA stem antibodies encode a Tyr in the CDR H3, the position of this Tyr varies, and some do not even have a Tyr in the CDR H3.^14^ Based on these observations, IGHV1-69 HA stem antibodies do not seem to have strong sequence preferences in the CDR H3 and light chain. At the same time, it is impossible that all IGHV1-69 antibodies can bind to HA stem, given that many IGHV1-69 antibodies are specific to other pathogens.^15^ Besides, some IGHV1-69 antibodies bind to HA head instead of HA stem.^16, 17^ Therefore, despite the first IGHV1-69 HA stem antibody being discovered 15 years ago,^9^ it remains unclear what sequence features make an IGHV1-69 antibody bind to the HA stem.

In this study, we performed two high-throughput experiments to probe the sequence constraints in the light chain and CDR H3 of the IGHV1-69 HA stem antibody CR9114, which is the broadest influenza neutralizing antibody known to date.^7^ Our first high-throughput experiment examined the compatibility of 78 light chain sequences from diverse IGHV1-69 antibodies with CR9114 heavy chain. Our findings indicated that despite having no contact with the HA stem epitope, light chain of CR9114 contained sequence determinants for its binding activity. Specifically, we demonstrated that the amino acid sequences at V_L_ residues 91 and 96 hugely influenced the light chain compatibility with CR9114 heavy chain for binding to HA stem. Of note, the Kabat numbering scheme is used throughout. Our second high-throughput experiment measured the binding affinity of 2,162 diverse CDR H3 variants to HA stem and showed that most CDR H3 variants, including many from other IGHV1-69 HA stem antibodies, were incompatible with CR9114. These results indicate that the sequence constraints in the CDR H3 and light chain of IGHV1-69 HA stem antibodies are stringent yet complex.

## RESULTS

### Light chain sequence of CR9114 is important for HA stem binding

A previous structural study has shown that CR9114, which is encoded by IGHV1-69 and IGLV1- 44, does not use light chain for binding (**Figure 1A**).^7^ In addition, the HA stem binding activity of CR9114 is not affected by replacing its light chain by one from an antibody of different specificity.^10^ Here, we aimed to systematically investigate if the sequence of CR9114 light chain is truly unimportant for HA stem binding. Briefly, we compiled a list of 120 antibodies from Genbank that are encoded by IGHV1-69 with different λ light chains (**Table S1**). The light chain sequences from these 120 antibodies were synthesized and paired with the germline sequence of CR9114 heavy chain to create a light chain variant library. We also included 10 light chain sequences with premature stop codons as negative controls. As a result, our light chain variant library contained 130 different light chain sequences, all of which paired with the germline sequence of CR9114 heavy chain. Of note, the germline sequence of CR9114 heavy chain was used because we wanted to avoid any incompatibility between heavy chain somatic hypermutations and light chain variants.

**Figure 1.**
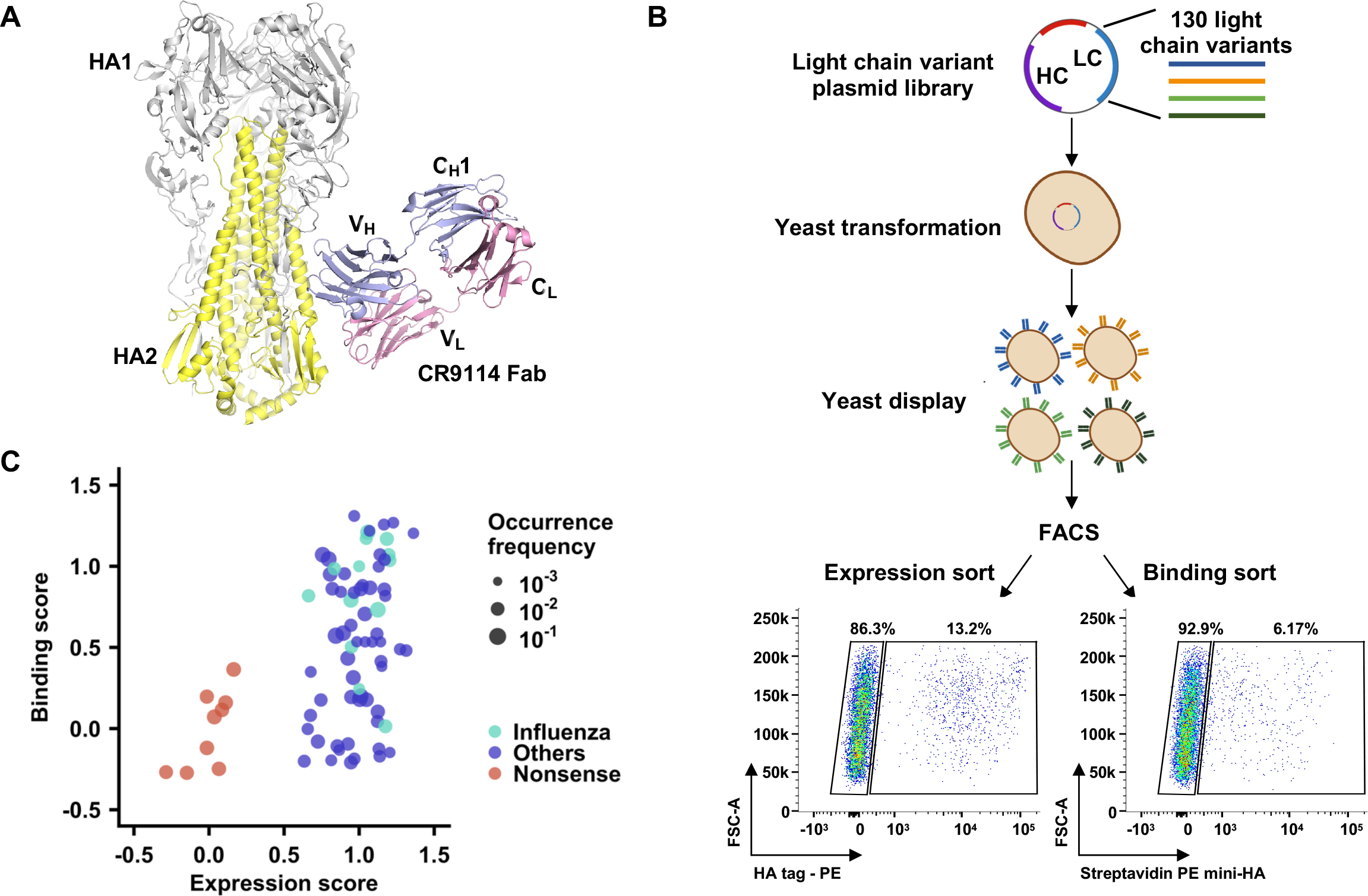
Yeast display of light chain variant library with CR9114 germline heavy chain. (A) Interaction between HA and CR9114 (PDB 4FQI).^7^ Grey: HA1; yellow: HA2; blue: heavy chain; pink: light chain. C_H_1 indicates constant region 1 of heavy chain and C_L_ indicates constant region 1 of light chain. V_H_ and V_L_ indicate variable regions of antibody heavy and light chains, respectively. **(B)** Schematic illustration of measuring the expression level and HA stem binding activity of many light chain variants in parallel using yeast surface display. Briefly, an antibody light chain (LC) variant library was paired with CR9114 germline heavy chain (HC), displayed on the yeast cell surface, and subjected to fluorescence-activated cell sorting (FACS) based on surface expression level and binding to mini-HA. The sorted populations were analyzed by next- generation sequencing. **(C)** Relationship between binding and expression scores. One data point represents one light chain variant. Cyan: light chain variants from known IGHV1-69 antibodies to influenza virus. Blue: light chain variants from IGHV1-69 antibodies to non- influenza antigens. Red: negative control variants with stop codons.

Subsequently, this light chain variant library was displayed on the yeast cell surface, and two- way sorted based on antibody expression level as well as binding activity to mini-HA, which is a trimeric HA stem construct without the head domain (**Figure 1B**).^5^ Four sorted populations were collected, namely no expression, high expression, no binding, and high binding. The frequency of each light chain variant in each sorted population was quantified by next-generation sequencing. Among the 130 light chain variants in the library, 78 had an average occurrence frequency of >0.2% across different sorted populations and were subjected to downstream analyses (**Table S1**). These 78 light chain variants include the one from CR9114 (i.e., wild type; WT), 13 from other influenza antibodies, 55 from non-influenza antibodies, and 9 negative controls with premature stop codons. For each light chain variant, an expression score and a binding score were computed based on its frequency in different sorted populations (**see Methods**). Both the expression and binding scores were normalized such that the scores for WT equaled 1 and the mean scores for negative controls (i.e., variants with stop codons) equaled 0. Pearson correlation coefficients of 0.75 and 0.74 were obtained between two replicates of the binding and expression sorts, respectively (**Figure S1**), confirming the reproducibility of our results.

Except for the negative controls, most light chain variants had an expression score of around 1 (**Figure 1C**), indicating that the light chain sequence of CR9114 germline had minimal influence on expression. However, many light chain variants had a low binding score (**Figure 1C**). In addition, the binding scores of light chain variants from influenza antibodies (mean = 0.84) were significantly higher than those from non-influenza antibodies (mean = 0.49, p-value = 0.005, two-tailed Student’s t-test), despite having similar expression scores (mean = 1.03 and 0.99, respectively, p-value = 0.35, two-tailed Student’s t-test). These results imply that the light chain sequence of CR9114 germline is an important determinant for its HA stem binding activity.

### Importance of CR9114 light chain residues 91 and 96

Next, we aimed to understand the molecular mechanism of how light chain modulated the HA stem binding activity of CR9114 germline. Since CR9114 light chain has no contact with the HA stem epitope (**Figure 1A**),^7^ its sequence determinants likely locates at the heavy-light chain interface. Structural analysis of the previously determined crystal structure of CR9114 in complex with HA^7^ suggested that V_L_ residues 91 and 96 in CDR L3 are critical for stabilizing the conformation of CDR H3, which in turn is important for binding (**Figure 2A**). CR9114 has an aromatic residue Trp at V_L_ residue 91, which forms an extensive π-π stacking network with four aromatic residues in CDR H3, namely V_H_ Y98, V_H_ Y99, V_H_ Y100, and V_H_ Y100a, as well as V_H_ W47 in the heavy chain framework region 2. In contrast, CR9114 has a small amino acid Ala at V_L_ residue 96, which points toward the heavy chain with limited space in between. These observations suggested that the compatibility of different light chain variants and CR9114 germline heavy chain depended on the amino acid identities at V_L_ residues 91 and 96. Specifically, we hypothesized that amino acids F/Y/W, which are both aromatic and bulky, were required at V_L_ residue 91 but forbidden at V_L_ residue 96.

**Figure 2.**
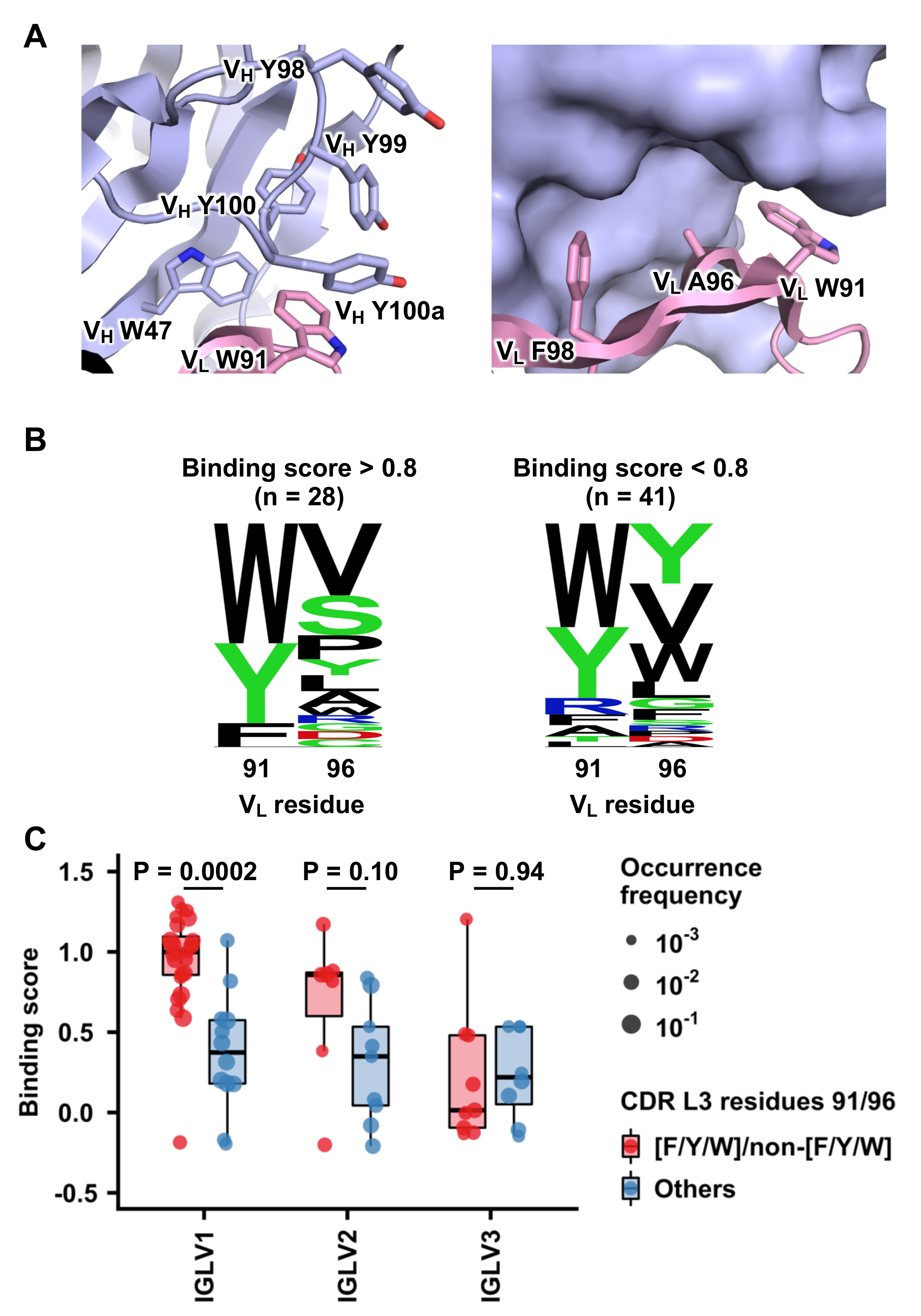
Light chain sequence determinants for HA stem binding activity of CR9114. (A) Left panel: an extensive π-π stacking network at the heavy-light chain interface of CR9114 that involves V_H_ W47, V_H_ Y98, V_H_ Y99, V_H_ Y100, V_H_ Y100a, and V_L_ W91. Right panel: Side chains of V_L_ W91, A96 and F98 at the heavy-light chain interface of CR9114, with heavy chain in surface representation. PDB 4FQI is used.^7^ Light blue: heavy chain; pink: light chain. V_H_ and V_L_ indicate variable regions of antibody heavy and light chains, respectively. **(B)** Sequence logos for V_L_ residues 91 and 96 of light chain variants with binding scores >0.8 (left panel) and <0.8 (right panel). Relative sizes of the letters indicate their frequency among the variants. **(C)** Binding scores of light chain variants in different IGLV families with and without V_L_ 91_F/Y/W_/96_non-F/Y/W_ are compared. Red: with V_L_ 91_F/Y/W_/96_non-F/Y/W_; Blue: without V_L_ 91_F/Y/W_/96_non-F/Y/W_. P-values were computed by two-tailed Student’s t-test.

We then compared the amino acid sequences at V_L_ residues 91 and 96 between high-binding and low-binding light chain variants, using an arbitrary binding score cutoff of 0.8 (**Figure 2B**). At V_L_ residue 91, all light chain variants with a binding score >0.8 contained an aromatic amino acid (i.e., W/Y/F), whereas non-aromatic amino acids could be observed among variants with a binding score <0.8. Conversely, aromatic amino acids were enriched at V_L_ residue 96 among variants with a binding score <0.8. These observations are consistent with our hypothesis above.

To further experimentally validate our findings, we introduced different mutations at residues 91 and 96 of CR9114 light chain, recombinantly expressed them by pairing with CR9114 germline heavy chain, and then tested their binding affinity to mini-HA. Hereafter, CR9114 with germline heavy chain is abbreviated as CR9114 gHC. The binding activity of CR9114 gHC to mini-HA was abolished by substituting V_L_ W91 with non-aromatic amino acids T/R/A, but not aromatic amino acids Y/F (**Table 1 and Figure S2**). Likewise, the binding activity of CR9114 gHC to mini-HA was abolished by substituting V_L_ A96 with aromatic amino acids W/F, but not non-aromatic amino acids V/S/R (**Table 1 and Figure S2**). Therefore, our binding experiment confirms that V_L_ residues 91 and 96 in the CDR L3 are important for CR9114 to interact with the HA stem, despite not being part of the paratope.

**Table 1.** Binding affinity of CR9114 gHC mutants against mini-HA. N.B. indicates no binding.

| **CR9114 gHC mutant Fab** | **K<sub>D</sub> (μM)** |
| --- | --- |
| WT | 6.2 |
| V<sub>L</sub> W91Y | 45.7 |
| V<sub>L</sub> W91F | 9.7 |
| V<sub>L</sub> W91T | N.B. |
| V<sub>L</sub> W91R | N.B. |
| V<sub>L</sub> W91A | N.B. |
| V<sub>L</sub> A96Y | N.B. |
| V<sub>L</sub> A96W | N.B. |
| V<sub>L</sub> A96F | N.B. |
| V<sub>L</sub> A96V | 3.8 |
| V<sub>L</sub> A96S | 1.9 |
| V<sub>L</sub> A96R | 13.0 |

### Additional sequence constraints in CR9114 light chain

Although our results indicated that aromatic amino acids were required at V_L_ residue 91 but forbidden at V_L_ residue 96 of CR9114, exceptions existed in our light chain variant library (**Table S1**). Specifically, we noticed that some light chain variants with a low binding score had aromatic (i.e., F/Y/W) and non-aromatic amino acids (i.e., non-F/Y/W) at V_L_ residues 91 and 96, respectively (i.e., V_L_ 91_F/Y/W_/96_non-F/Y/W_). This observation suggests that there are additional sequence constraints in the light chain of CR9114.

Members of our light chain variant library were from three IGLV families, namely IGLV1, IGLV2, and IGLV3. For light chain variants in IGLV1 family, those with V_L_ 91_F/Y/W_/96_non-F/Y/W_ had significantly higher binding scores than those without (p-value = 2e-4, two-tailed Student’s t-test, **Figure 2C**). In contrast, while the light chain variants in IGLV2 family with V_L_ 91_F/Y/W_/96_non-F/Y/W_ also had higher binding scores than those without, such difference was not as significant (p-value = 0.10, two-tailed Student’s t-test, **Figure 2C**). Furthermore, the binding scores of light chain variants in IGLV3 family with and without the V_L_ 91_F/Y/W_/96_non-F/Y/W_ motif had no significant difference (p-value = 0.94, two-tailed Student’s t-test) and were generally low (**Figure 2C**). Of note, in all three IGLV families, the expression scores of light chain variants with and without V_L_ 91_F/Y/W_/96_non-F/Y/W_ had no significant difference (p-values ranging from 0.31 to 0.84, two-tailed Student’s t-test, **Figure S3A**). These results show that besides the amino acid sequences at V_L_ residues 91 and 96, sequence differences among IGLV families could also influence the binding activity of CR9114 to HA stem.

Our additional analysis suggested that a CDR L3 length of 11 was optimal for CR9114 binding to the HA stem (**Figure S3B**), since the binding scores of light chain variants lowered when the CDR L3 lengths deviated from 11. In contrast, the expression scores were similar across light chain variants with different CDR L3 lengths (**Figure S3C**). We also analyzed the relationship between binding scores and the number of V_L_ somatic hypermutations (SHMs), but no significant correlation was found (p-values ranging from 0.39 to 0.98, Pearson correlation test, **Figure S3D-F**). Overall, our analyses demonstrate that light chain germline features, including CDR L3 length and germline gene usage, are important sequence determinants for the HA stem binding activity of CR9114.

### High-throughput characterization of CDR H3 variants

Similar to the light chain, the sequence constraints of the CDR H3 of IGHV1-69 HA stem antibodies have also been unclear, especially since they are highly diverse.^14^ Therefore, we next aimed to probe the sequence constraints of CR9114 CDR H3. Briefly, we compiled a list of 3,325 CDR H3 sequences from diverse IGHV1-69 antibodies from Genbank (**Table S2**). Additionally, we performed a site-saturation mutagenesis of the CDR H3 of CR9114. Together with the WT CDR H3 sequence of CR9114, a CDR H3 library with 3,606 variants in CR9114 gHC was constructed and displayed on yeast cell surface. Tite-Seq^18^ was then applied to measure the apparent dissociation constant (K_D_) values of individual variants to mini-HA (**Figure 3A**). At the same time, antibody expression level was measured by two-way cell sorting and next-generation sequencing, as described above for the light chain variant library. The expression score for each CDR H3 variant was normalized such that the score for WT equaled 1 and the mean score for nonsense mutants of CR9114 CDR H3 equaled 0.

**Figure 3.**
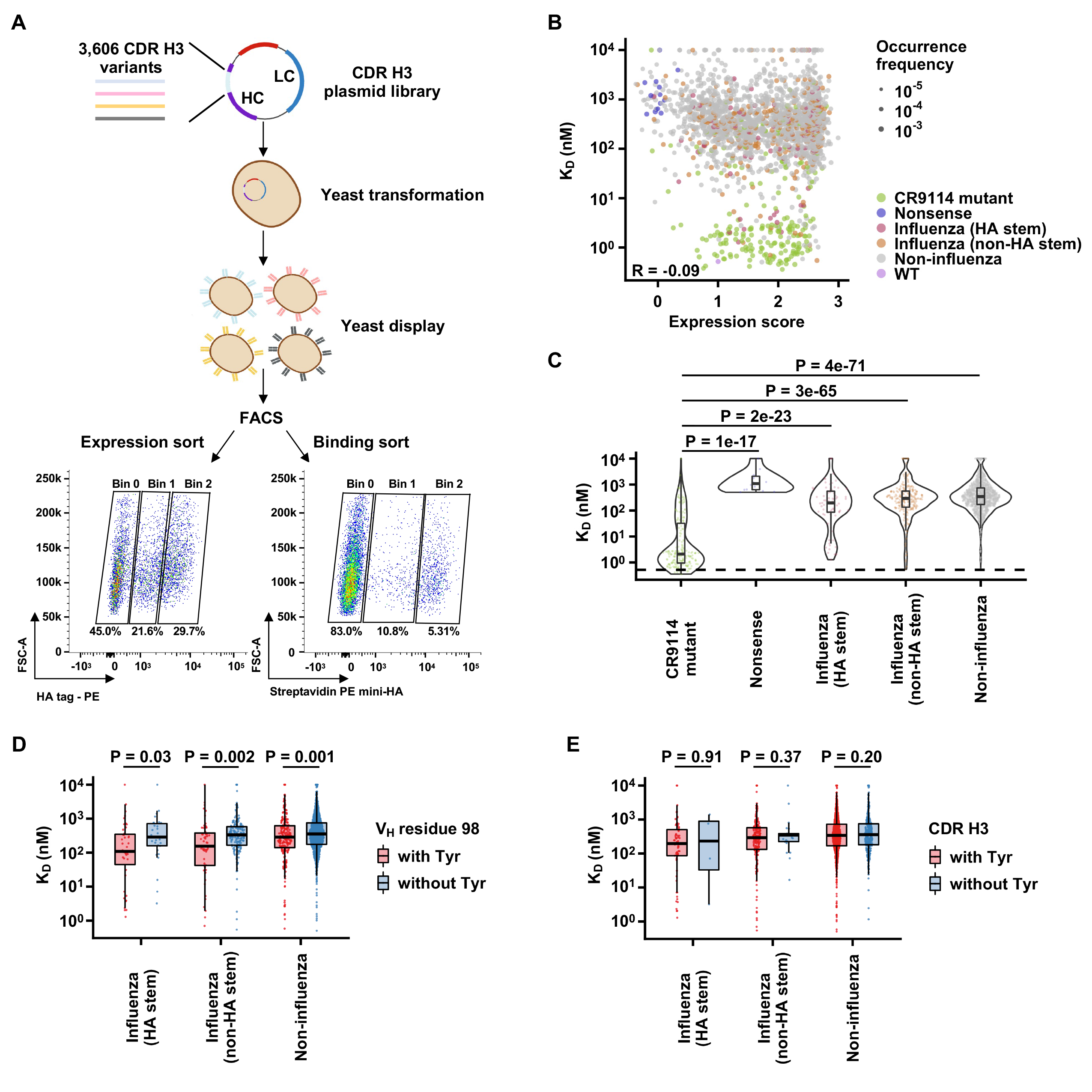
Yeast display of CDR H3 library of CR9114 gHC. (A) Schematic illustration of measuring the expression level and HA stem binding activity of many CDR H3 variants in parallel using yeast surface display. Briefly, a CDR H3 library of CR9114 gHC was displayed on the yeast cell surface and subjected to FACS based on surface expression level and binding to mini-HA. The sorted populations were analyzed by next-generation sequencing. **(B)** Relationship between apparent dissociation constant (K_D_) values and expression scores. Greenish yellow: CR9114 single amino acid mutants; purple: nonsense variants with stop codons; red: CDR H3 variants from IGHV1-69 HA stem antibodies; orange: CDR H3 variants from IGHV1-69 non-HA stem influenza antibodies; grey: CDR H3 variants from IGHV1-69 non- influenza antibodies; pink: WT. The Pearson correlation coefficient (R) is indicated. **(C)** Distribution of the apparent K_D_ values among CDR H3 variants from different types of antibodies. **(D)** Comparison of apparent K_D_ values between antibodies with and without V_H_ Y98. **(E)** Comparison of apparent K_D_ values between antibodies with and without a Tyr in their CDR H3. P-values were computed by two-tailed Student’s t-test.

After filtering out CDR H3 variants with low occurrence frequency or noisy estimation in the apparent K_D_ values (**see Methods**), 2,162 CDR H3 variants were subjected to downstream analyses (**Table S2**). These included the WT CDR H3 of CR9114, 206 single amino acid mutants of CR9114 CDR H3, 14 nonsense mutants of CR9114 CDR H3, 73 CDR H3 variants from HA stem antibodies, 245 from non-HA stem influenza antibodies, and 1,623 from non-influenza antibodies. Pearson correlations of 0.71 and 0.61 were obtained between two replicates of Tite-Seq and expression sort, respectively (**Figure S4A-B**), confirming the reproducibility of our results. As expected, the expression score of nonsense mutants of CR9114 CDR H3 was significantly lower than other CDR H3 variants (p-values = 2e-22 to 1e-47, two-tailed Student’s t-test, **Figure S4D**). Similarly, the binding affinity of nonsense mutants of CR9114 CDR H3 to mini-HA was significantly weaker than that of single amino acid mutants of CR9114 CDR H3 (p-value = 1e-17, two-tailed Student’s t-test, **Figure 3B-C**). These results validated the quality of our data.

### Most CDR H3 variants are incompatible with CR9114 for HA stem binding

While the apparent K_D_ values of many single amino acid mutants of CR9114 CDR H3 to mini-HA were around 1 nM, those of CDR H3 variants from other antibodies, including HA stem antibodies, were mostly between 100 nM to 1 μM (**Figure 3B-C**). In contrast, the expression scores of single amino acid mutants of CR9114 CDR H3 and other CDR H3 variants were similar (p-values = 0.22 to 0.76, two-tailed Student’s t-test, **Figure S4D**). Of note, several CDR H3 variants from non-influenza antibodies had an apparent K_D_ of around 1 nM (**Figure S4C and Table S2**). However, when we recombinantly expressed one of these CDR H3 variants, it did not show binding to mini-HA, indicating there were false positives in our Tite-Seq experiment (**Figure S4E**). Together, these observations suggest that the CDR H3 of CR9114 has a stringent sequence requirement for HA stem binding.

A previous study has demonstrated that many IGHV1-69 HA stem antibodies, including CR9114, are featured by a CDR H3-encoded Tyr at V_H_ residue 98 that interacts extensively with the HA stem epitope.^14^ Consistently, our results showed that CDR H3 variants with Y98 had slightly, yet significantly, better apparent K_D_ values (p-values = 0.001 to 0.03, two-tailed Student’s t-test, **Figure 3D**). In contrast, CDR H3 variants with and without Tyr, regardless of the residue position, did not show any significant difference in apparent K_D_ values (p-values = 0.20 to 0.91, two-tailed Student’s t-test, **Figure 3E**). Therefore, our result substantiates that V_H_ Y98 partially contributes to the compatibility of CDR H3 variants with CR9114 for HA stem binding. However, given that many CDR H3 variants from IGHV1-69 HA stem antibodies with Y98 are incompatible with CR9114 gHC for binding (**Figure 3D**), the sequence constraints of CR9114 CDR H3 likely involve other CDR H3 residues.

### Importance of non-paratope residues in CR9114 CDR H3

To further understand the sequence constraints of CR9114 CDR H3, we aimed to identify residues that are key for HA stem binding. Subsequently, we analyzed the single amino acid mutants of CR9114 CDR H3 in our CDR H3 library. Our Tite-Seq result indicated that most mutations at the four consecutive Tyr residues in the CDR H3, namely V_H_ Y98, V_H_ Y99, V_H_ Y100, and V_H_ Y100a, weakened the binding affinity of CR9114 gHC (**Figure 4A**). This observation is consistent with our structural analysis above (**Figure 2A**), showing that these four Tyr residues form an extensive π-π stacking network with V_H_ W47 and V_L_ W91, which is essential for the conformational stability of the CDR H3. Additionally, our Tite-Seq result also revealed the low mutational tolerance of V_H_ H95 and V_H_ N97, hence their importance in the HA stem binding activity of CR9114 (**Figure 4A**).

**Figure 4.**
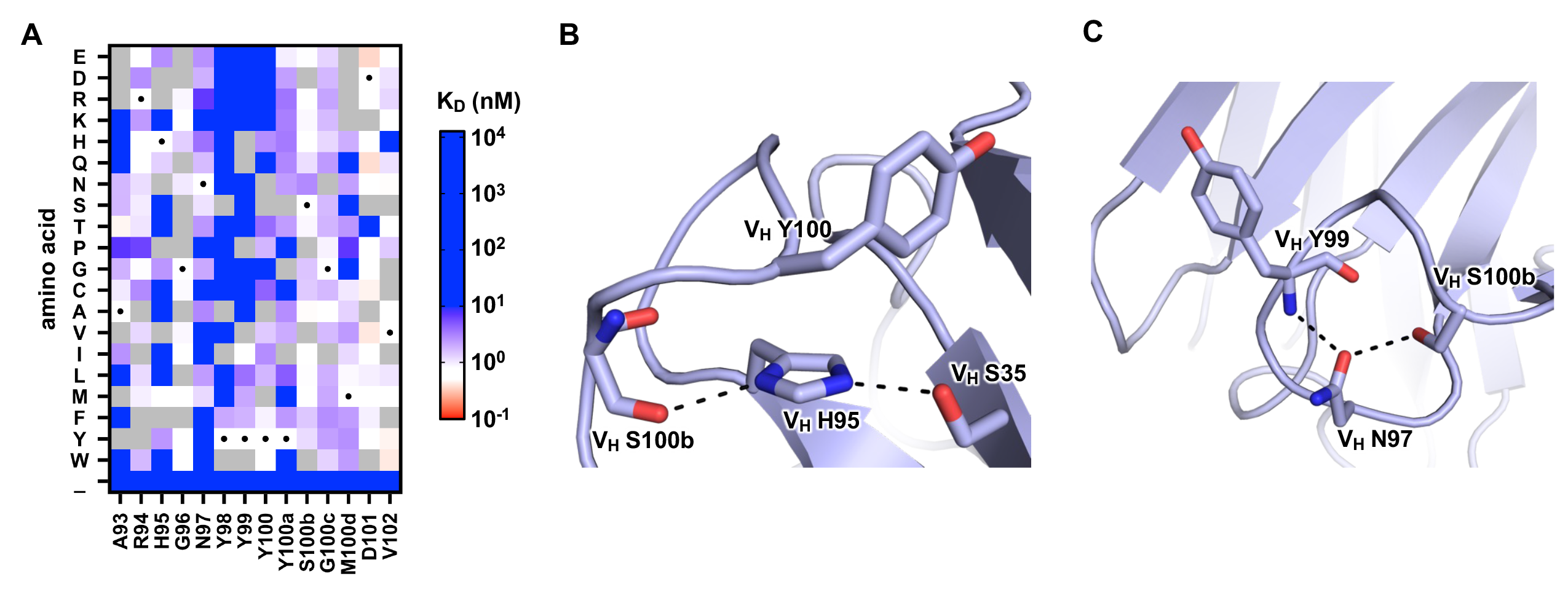
Importance of V_H_ H95 and N97 for HA stem binding activity of CR9114. (A) Apparent K_D_ values of individual mutations in the CDR H3 of CR9114 are shown as a heatmap. Wild-type (WT) amino acids are indicated by a black circle. **(B)** Intramolecular interactions involving CR9114 V_H_ H95 are shown. **(C)** Intramolecular Interactions involving CR9114 V_H_ N97 are shown. Hydrogen bonds are represented by dashed lines. PDB 4FQI is used.^7^

Based on a previously determined crystal structure of CR9114 in complex with HA,^7^ the side chains of both V_H_ H95 and V_H_ N97 are not interacting with the HA stem epitope. Instead, both V_H_ H95 and V_H_ N97 form intramolecular interactions to stabilize the CDR H3 conformation. V_H_ H95 H-bonds with V_H_ S100b and V_H_ S35 as well as interacts with V_H_ Y100 via T-shaped π-π stacking (**Figure 4B**), whereas V_H_ N97 H-bonds with V_H_ Y99 and V_H_ S100b (**Figure 4C**). These observations corroborate our light chain analysis, demonstrating that non-paratope residues that stabilize the CDR H3 conformation are important for the HA stem binding activity of CR9114.

## DISCUSSION

IGHV1-69 is one of the most highly used heavy chain V genes in the human antibody repertoire,^19^ suggesting its importance in the immune system. In fact, many known broadly neutralizing antibodies to various pathogens, such as influenza virus, hepatitis C virus (HCV), and human immunodeficiency virus (HIV), were encoded by IGHV1-69.^15^ As a result, IGHV1-69 is regarded as an S.O.S. component of the human antibody repertoire.^20^ However, sequence determinants for the antigen specificity of IGHV1-69 antibodies have been largely elusive. In this study, we used a high-throughput approach to probe the sequence determinants for the HA stem binding activity of CR9114, an IGHV1-69 broadly neutralizing antibody to influenza HA stem.^7^ Our results revealed the importance of the CR9114 light chain in binding, albeit no contact with the epitope. In addition, we showed that the CDR H3 sequence of CR9114 has stringent sequence constraints. Overall, this work advances our understanding of the sequence determinants that define IGHV1-69 HA stem antibodies.

Our results show that V_L_ residue 96 in CDR L3 is a determinant for the HA stem binding activity of CR9114. V_L_ residue 96 in some IGHV1-69 antibodies, including CR9114, is encoded by light chain J gene (**Figure S5A**). Among seven known IGLJ germline genes, three (IGLJ1, IGLJ4, and IGLJ5) encode an aromatic amino acid at V_L_ residue 96 (**Figure S5B**). Given that aromatic amino acids are forbidden at V_L_ residue 96 of CR9114, these observations suggest that light chain J gene plays a role in generating IGHV1-69 HA stem antibodies. Of note, the contribution of light chain J gene to antibody binding is rarely reported, if any, since most antibody studies focus on the V genes. However, the sequence determinants for the binding activity of IGHV1-69 HA stem antibodies likely vary from antibody to antibody (see discussion below). As a result, additional work is needed to decipher whether light chain J gene usage is a common sequence constraint of IGHV1-69 HA stem antibodies.

Perhaps the most perplexing result in our study is that most CDR H3 sequences from other IGHV1-69 HA stem antibodies are incompatible with CR9114. Provided that diverse light chain sequences can be observed in IGHV1-69 HA stem antibodies,^7, 9, 11–13^ the compatibility of a given CDR H3 sequence in IGHV1-69 HA stem antibodies likely depends on their light chain sequences. In other words, we postulate that whether an IGHV1-69 antibody can bind to HA stem is coordinately determined by the light chain and CDR H3 sequences. Consistently, our results revealed the importance of light chain-CDR H3 interaction in the HA stem binding activity of CR9114. Therefore, the sequence determinants of IGHV1-69 HA stem antibodies are stringent yet complex, which explain the lack of sequence convergence among IGHV1-69 HA stem antibodies. Comprehending these sequence determinants will enable an accurate estimation of the proportion of B cells that can give rise to IGHV1-69 HA stem antibodies, especially those that can cross-react to both group 1 and 2 HAs as well as influenza B HA, like CR9114. As the efforts to develop a universal influenza vaccine continue,^21–23^ future studies on the sequence determinants of IGHV1-69 HA stem antibodies are warranted.

## Limitations of the study

Since our approach has focused on λ light chain of IGHV1-69 antibodies, we were unable to demonstrate whether the κ light chain has similar sequence constraints when paired with CR9114 heavy chain for HA stem binding. Another limitation of our study is that only five antigen concentrations were used in our Tite-Seq experiment as opposed >10 in other Tite-Seq studies.^18, 24, 25^ This shortcoming would increase the estimation error of apparent K_D_ values.

## ACKNOWLEDGEMENTS

This work was supported by the National Institutes of Health R01 AI167910 (N.C.W.), DP2 AT011966 (N.C.W.), the Searle Scholars Program (N.C.W.), and Howard Hughes Medical Institute Emerging Pathogens Initiative (N.C.W.).

## AUTHOR CONTRIBUTIONS

N.C.W. conceived and designed the study. Y.W., H.L., and N.C.W. assembled the dataset.

Q.W.T. constructed the yeast libraries and prepared the sequencing libraries. Y.W., and N.C.W. performed data analysis. H.L., T.J.C.T., R.L., and K.J.M. assisted with experiments. Q.W.T., Y.W., H.L., and N.C.W. wrote the paper and all authors reviewed and/or edited the paper.

## DECLARATION OF INTERESTS

N.C.W. consults for HeliXon. The authors declare no other competing interests.

## METHODS

### CR9114 gHC yeast display plasmid

CR9114 gHC yeast display plasmid, pCTcon2_CR9114_GL, was generated by cloning the coding sequences of (from N-terminal to C-terminal, all in-frame) Aga2 secretion signal, CR9114 wild-type Fab light chain, V5 tag, equine rhinitis B virus (ERBV-1) 2A self-cleaving peptide, Aga2 secretion signal, CR9114 germline Fab heavy chain, HA tag, and Aga2p, into the pCTcon2 vector.^26^

### Construction of CDR H3 library and light chain variant library

Sequences of IGHV1-69 antibodies were obtained from GenBank (https://www.ncbi.nlm.nih.gov/genbank/).27 IgBLAST was used to identify the CDR H3 region.^28^ Light chain variant library (**Table S1**) and CDR H3 library (**Table S2**) were synthesized as oligo pools by Integrated DNA Technologies and Twist Bioscience, respectively. Names and sequences of primers for cloning the libraries into pCTcon2_CR9114_GL are listed in **Table S3**. Oligo pool of the light chain variant library was PCR-amplified using primers IGHV1-69- Lightchain-lib-IF and IGHV1-69-Lightchain-lib-IR. Then, the amplified oligonucleotide pool was gel-purified using a Monarch DNA Gel Extraction Kit (NEB). To generate the linearized vector for the light chain variant library, pCTcon2_CR9114_GL was used as a template for PCR using primers IGHV1-69-Lightchain-lib-VF and IGHV1-69-Lightchain-lib-VR. The PCR product was then gel-purified. Similarly, oligo pool of the CDR H3 library was PCR-amplified using primers IGHV1-69-CDRH3-lib-IF and IGHV1-69-CDRH3-lib-IR. Then, the amplified oligo pool was gel-purified using a Monarch DNA Gel Extraction Kit (NEB). To generate the linearized vector for the light chain variant library, pCTcon2_ CR9114_GL was used as a template for PCR using primers IGHV1-69-CDRH3-lib-VF and IGHV1-69-CDRH3-lib-VR. The PCR product was then gel-purified. All PCRs were performed using KOD Hot Start DNA polymerase (EMD Millipore) according to the manufacturer’s instructions.

### Yeast transformation

Yeast cells were transformed by electroporation following a previously described protocol.^29^ Briefly, *Saccharomyces cerevisiae* EBY100 cells (American Type Culture Collection) were grown in YPD medium (1% w/v yeast nitrogen base, 2% w/v peptone, 2% w/v D(+)- glucose) overnight at 30::J°C with shaking at 225::Jrpm until OD_600_ reached 3. Then, an aliquot of overnight culture was grown in 100::JmL YPD media, with an initial OD_600_ of 0.3, shaking at 225::Jrpm at 30::J°C. Once OD_600_ reached 1.6, yeast cells were collected by centrifugation at 1700::J×::J*g* for 3::Jmin at room temperature. Media were removed and the cell pellet was washed twice with 50::JmL ice-cold water, and then once with 50::JmL of ice-cold electroporation buffer (1::JM sorbitol, 1::JmM calcium chloride). Cells were resuspended in 20::JmL conditioning media (0.1::JM lithium acetate, 10::JmM dithiothreitol) and shaked at 225::Jrpm at 30::J°C. Cells were collected via centrifugation at 1700::J×::J*g* for 3::Jmin at room temperature, washed once with 50::JmL ice-cold electroporation buffer, resuspended in electroporation buffer to reach a final volume of 1::JmL, and kept on ice. 5::Jµg of the amplified oligo pool (light chain variant library or CDR H3 library) and 4::Jµg of the corresponding purified linearized vector were added into 400::JµL of conditioned yeast. The mixture was transferred to a pre-chilled BioRad GenePulser cuvette with 2::Jmm electrode gap and kept on ice for 5::Jmin until electroporation. Cells were electroporated at 2.5::JkV and 25::JµF, achieving a time constant between 3.7 and 4.1::Jms. Electroporated cells were transferred into 4::JmL of YPD media supplemented with 4::JmL of 1::JM sorbitol and incubated at 30::J°C with shaking at 225::Jrpm for 1::Jh. Cells were collected via centrifugation at 1700::J×::J*g* for 3::Jmin at room temperature, resuspended in 0.6::JmL SD-CAA medium (2% w/v D-glucose, 0.67% w/v yeast nitrogen base with ammonium sulfate, 0.5% w/v casamino acids, 0.54% w/v Na_2_HPO_4_, 0.86% w/v NaH_2_PO_4_·H_2_O, all dissolved in deionized water), plated onto SD-CAA plates (2% w/v D-glucose, 0.67% w/v yeast nitrogen base with ammonium sulfate, 0.5% w/v casamino acids, 0.54% w/v Na_2_HPO_4_, 0.86% w/v NaH_2_PO_4_·H_2_O, 18.2% w/v sorbitol, 1.5% w/v agar, all dissolved in deionized water) and incubated at 30::J°C for 40::Jh. Colonies were then collected in SD-CAA medium, centrifuged at 1700::J×::J*g* for 5::Jmin at room temperature, and resuspended in SD-CAA medium with 15% v/v glycerol such that OD_600_ was 50. Glycerol stocks were stored at -80::J°C until used.

### Expression and purification of mini-HA

The mini-HA #4900 protein^5^ was fused with N-terminal gp67 signal peptide and a C-terminal BirA biotinylation site, thrombin cleavage site, trimerization domain, and a His_6_ tag, and then cloned into a customized baculovirus transfer vector^30^. Recombinant bacmid DNA that carried the mini-HA domain was generated using the Bac-to-Bac system (Thermo Fisher Scientific) according to the manufacturer’s instructions. Baculovirus was generated by transfecting the purified bacmid DNA into adherent Sf9 cells using Cellfectin reagent (Thermo Fisher Scientific) according to the manufacturer’s instructions. The baculovirus was further amplified by passaging in adherent Sf9 cells at a multiplicity of infection (MOI) of 1. Recombinant mini-HA protein was expressed by infecting 1::JL of suspension Sf9 cells at an MOI of 1. On day 3 post-infection, Sf9 cells were pelleted by centrifugation at 4000::J×::J*g* for 25::Jmin, and soluble recombinant mini-HA was purified from the supernatant by affinity chromatography using Ni Sepharose excel resin (Cytiva) and then size exclusion chromatography using a HiLoad 16/100 Superdex 200 prep grade column (Cytiva) in 20::JmM Tris-HCl pH 8.0, 100::JmM NaCl. The purified mini-HA protein was concentrated by Amicon spin filter (Millipore Sigma) and filtered by 0.22 µm centrifuge Tube Filters (Costar). Concentration of the protein was determined by nanodrop (Fisher Scientific). Protein was subsequent aliquoted, flash frozen by dry-ice ethanol mixture, and store at -80°C until used.

### Biotinylation and PE-conjugation of mini-HA

Purified mini-HA was biotinylated using the Biotin-Protein Ligase-BIRA kit according to the manufacturer’s instructions (Avidity). Biotinylated mini-HA was then conjugated to streptavidin– PE (Thermo Fisher Scientific) by incubating at room temperature for 15 mins.

### Fluorescence-activated cell sorting (FACS) of yeast display library

100::JµL glycerol stock of the yeast display library was recovered in 50 mL SD-CAA medium by incubating at 27::J°C with shaking at 250::Jrpm until OD_600_ reached between 1.5 and 2.0. Then 15::JmL of the yeast culture was harvested and pelleted via centrifugation at 4000::J×::J*g* at 4::J°C for 5::Jmin. The supernatant was discarded, and SGR-CAA (2% w/v galactose, 2% w/v raffinose, 0.1% w/v D-glucose, 0.67% w/v yeast nitrogen base with ammonium sulfate, 0.5% w/v casamino acids, 0.54% w/v Na_2_HPO_4_, 0.86% w/v NaH_2_PO_4_·H_2_O, all dissolved in deionized water) was added to make up the volume to 50::JmL. The yeast culture was then transferred to a baffled flask and incubated at 18::J°C with shaking at 250::Jrpm. Once OD_600_ reached between 1.3 and 1.6, 1::JmL of yeast culture was harvested and pelleted via centrifugation at 4000::J×::J*g* at 4::J°C for 5::Jmin. The pellet was subsequently washed with 1::JmL of 1× PBS twice. After the final wash, cells were resuspended in 1::JmL of 1× PBS.

For expression sort, PE anti-HA.11 (epitope 16B12, BioLegend, Cat. No. 901517) that was buffer-exchanged into 1× PBS was added to the cells at a final concentration of 1::Jµg/mL. For binding sort of the light chain variant library, PE-conjugated mini-HA was added to washed cells at a final concentration of 30::JnM. For Tite-Seq, cells were labeled with PE-conjugated mini-HA at each of five antigen concentrations (one-log increments spanning 0.003 nM to 30 nM). A negative control was set up with nothing added to the PBS-resuspended cells. Samples were incubated overnight at 4::J°C with rotation. Then, the yeast pellet was washed twice in 1× PBS and resuspended in FACS tubes containing 2::JmL 1× PBS. Using a BD FACS Aria II cell sorter (BD Biosciences) and FACS Diva software v8.0.1 (BD Biosciences), cells in the selected gates were collected in 1::JmL of SD-CAA containing 1× penicillin/streptomycin. Single yeast cells were gated by forward scatter (FSC) and side scatter (SSC). Single cells were then gated by PE anti-HA.11 for expression sort. For Tite-Seq, single cells were gated into three bins along the PE-A axis based on unstained and CR9114 gHC controls, with bin 0 comprising all PE negative cells, bin 2 comprising PE positive cells with comparable expression or binding affinity to the germline CR9114 positive population, and bin 1 comprising the intermediate population between bin 0 and bin 2. Cells were then collected via centrifugation at 3800::J×::J*g* at 20::J°C for 15::Jmin. The supernatant was discarded. Subsequently, the pellet was resuspended in 100::JµL of SD- CAA and plated on SD-CAA plates at 30::J°C. After 40::Jh, colonies were collected in 2::JmL of SD-CAA. Frozen stocks were made by reconstituting the pellet in 15% v/v glycerol (in SD-CAA medium) and then stored at -80::J°C until used. FlowJo v10.8 software (BD Life Sciences) was used to analyze FACS data.

### Next-generation sequencing of light chain variant library and CDR H3 library

Plasmids from the yeast cells were extracted using a Zymoprep Yeast Plasmid Miniprep II Kit (Zymo Research) following the manufacturer’s protocol. The CDR H3 library was subsequently amplified by PCR using primers IGHV1-69-CDRH3-recover-F and IGHV1-69-CDRH3-recover-R whereas the light chain variant library was amplified using primers IGHV1-69-Lightchain- recover-F and IGHV1-69-Lightchain-R. Subsequently, adapters containing sequencing barcodes were appended to the amplicon using primers 5′-AAT GAT ACG GCG ACC ACC GAG ATC TAC ACX XXX XXX XAC ACT CTT TCC CTA CAC GAC GCT-3′, and 5′-CAA GCA GAA GAC GGC ATA CGA GAT XXX XXX XXG TGA CTG GAG TTC AGA CGT GTG CT-3′.

Positions annotated by an “X” represented the nucleotides for the index sequence. All PCRs were performed using Q5 High-Fidelity DNA polymerase (NEB) according to the manufacturer’s instructions. PCR products were purified using PureLink PCR Purification Kit (Thermo Fisher Scientific). The final PCR products were submitted for next generation sequencing using NovaSeq SP PE250 (Illumina).

### Computing the binding score, expression score, and apparent K_D_ values

The sequencing data was initially obtained in FASTQ format and subsequently analyzed using a custom python Snakemake pipeline.^31^ Briefly, PEAR was used for merging the forward and reverse reads.^32^ The number of reads corresponding to each variant in each sample is counted.

A pseudocount of 1 was added to the final count to avoid division by zero in downstream analysis. The binding and expression enrichment values of each variant *var* were computed as follows:

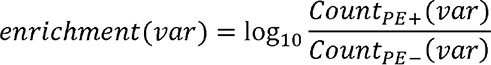

where the *Count*_*PE*+_(*var*) is the read count of variant *var* in the PE positive sample for a given binding or expression sort, while *Count*_*PE*−_(*var*) is the read count of variant *var* in the PE negative sample. The binding and expression scores for each variant *var* were further computed from the enrichment values as follows:

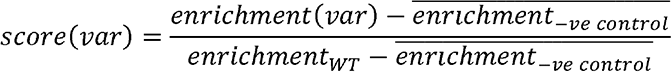

where 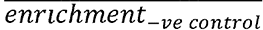 is the average enrichment value of the negative controls with stop codons and *enrichment*_*WT*_ is the enrichment value of the WT. The final score for each variant *var* is the average of two biological replicates.

To compute apparent K_D_ value of each CDR H3 variant from the Tite-Seq data, we adopted the analysis approach as previously described.^33^ Briefly, to determine the mean bin of PE fluorescence for each CDR H3 variant *var* at each mini-HA concentration, a simple weighted mean calculation was applied:

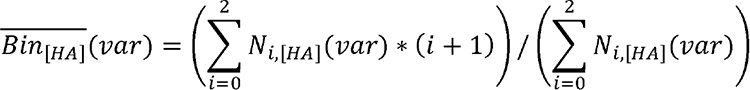

where N_i,[HA]_(*var*) is the number of cells with CDR H3 variant *var* that fall into bin *i* at mini-HA concentration [*HA*]. This calculation computes a weighted average by assigning integer weight to the bin *i*.

For each CDR H3 variant *var* we estimated its sorted cell count N_i,[HA]_(*var*) that corresponds to bin i at mini-HA concentration [*HA*] as follows:

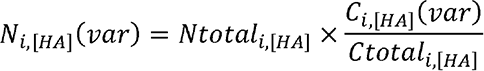

where variant read count *C*_i,[HA]_(*var*) is the read counts for CDR H3 variant *var* in bin *i* at mini- HA concentration [*HA*], *Ctotal*_*i,[HA]*_ is the total read counts for bin *i* at mini-HA concentration [*HA*], *Ntotal*_*i,[HA]*_ is the total number of cells in bin i at mini-HA concentration [*HA*].

We then determined the apparent K_D_ value for each variant K_D_(*var*) HA via a nonlinear least-squares regression using a standard non-cooperative Hill equation:

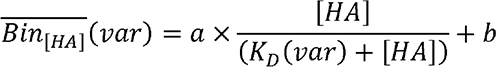

where free parameters *a* is titration response range and *b* is titration curve baseline.

CDR H3 variants with a total cell counts of less than 30 across bins 0 to 2 were discarded from our analysis. CDR H3 variants with an apparent KD value <10^-8^ and a p-value >0.2 were also discarded. Light chain variants with an average occurrence frequency ≤0.2% across different sorted populations were discarded from our analysis. Sequence logos were generated by Logomaker in Python.^34^

### Expression and purification of Fabs

The heavy and light chains of Fab were cloned into phCMV3 vector. Light chain mutants were generated using the QuikChange XL Mutagenesis kit (Stratagene) following the manufacturer’s instructions. The plasmids were co-transfected into Expi293F cells at a 2:1 (HC:LC) mass ratio using ExpiFectamine 293 Reagent (Thermo Fisher Scientific). At 7 days post-transfection, the supernatant was collected, and the Fab was purified using a CaptureSelect CH1-XL Pre-packed Column (Thermo Fisher Scientific).

### Biolayer interferometry binding assay

Binding assays were performed by biolayer interferometry (BLI) using an Octet Red96e instrument (FortéBio) at room temperature as described previously.^35^ Briefly, His-tagged mini-HA proteins at 0.5::JμM in 1× kinetics buffer (1× PBS, pH 7.4, 0.01% w/v BSA and 0.002% v/v Tween 20) were loaded onto anti-Penta-HIS (HIS1K) biosensors and incubated with the indicated concentrations of Fabs. The assay consisted of five steps: (1) baseline: 60::Js with 1× kinetics buffer; (2) loading: 60::Js with His-tagged mini-HA proteins; (3) baseline: 60::Js with 1× kinetics buffer; (4) association: 60::Js with Fab samples; and (5) dissociation: 60::Js with 1× kinetics buffer. For estimating the exact *K*_D_, a 1:1 binding model was used.

### Data availability

Raw sequencing data have been submitted to the NIH Short Read Archive under BioProject: PRJNA976657.

### Code availability

Custom python scripts for all analyses have been deposited to: https://github.com/nicwulab/CR9114_LC_CDRH3_screen

---

**Supplementary Figure 1.**
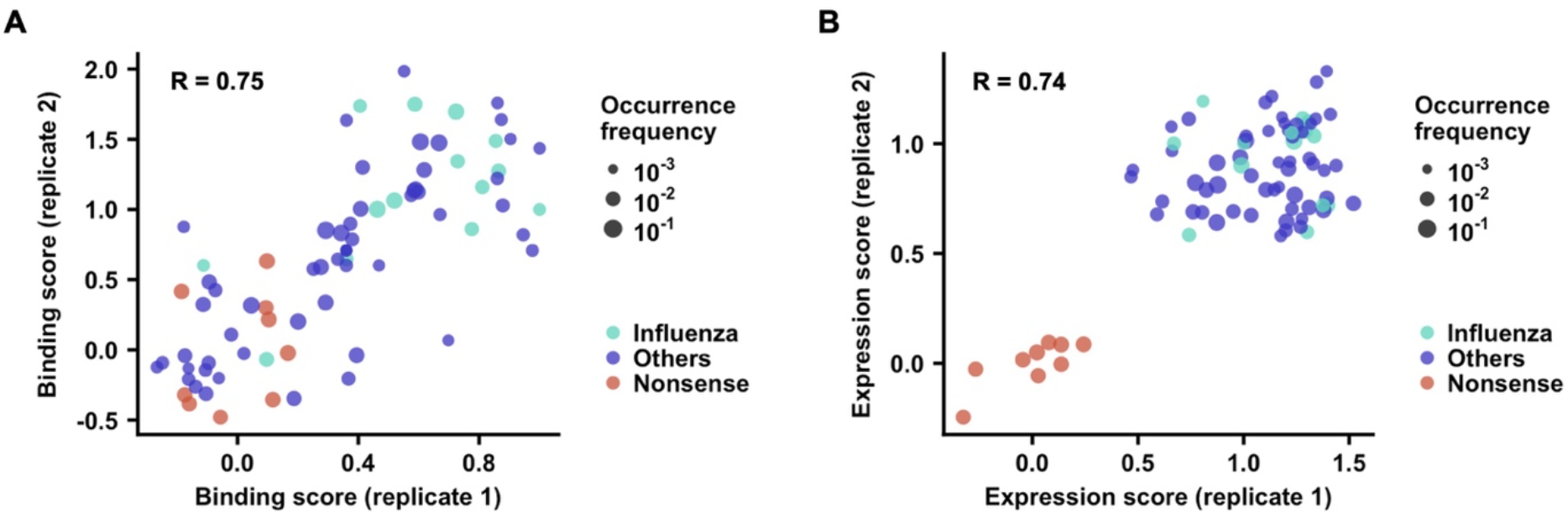
Correlation between biological replicates of binding and expression sorts of CR9114 light chain variant library. (A) Correlation of binding scores between two independent biological replicates is shown. (B) Correlation of expression scores between two independent biological replicates is shown. Cyan: light chain variants from known IGHV1-69 antibodies to influenza virus. Blue: light chain variants from IGHV1-69 antibodies to non-influenza antigens. Red: negative control variants with premature stop codons. The Pearson correlation coefficient (R) is indicated.

**Supplementary Figure 2.**
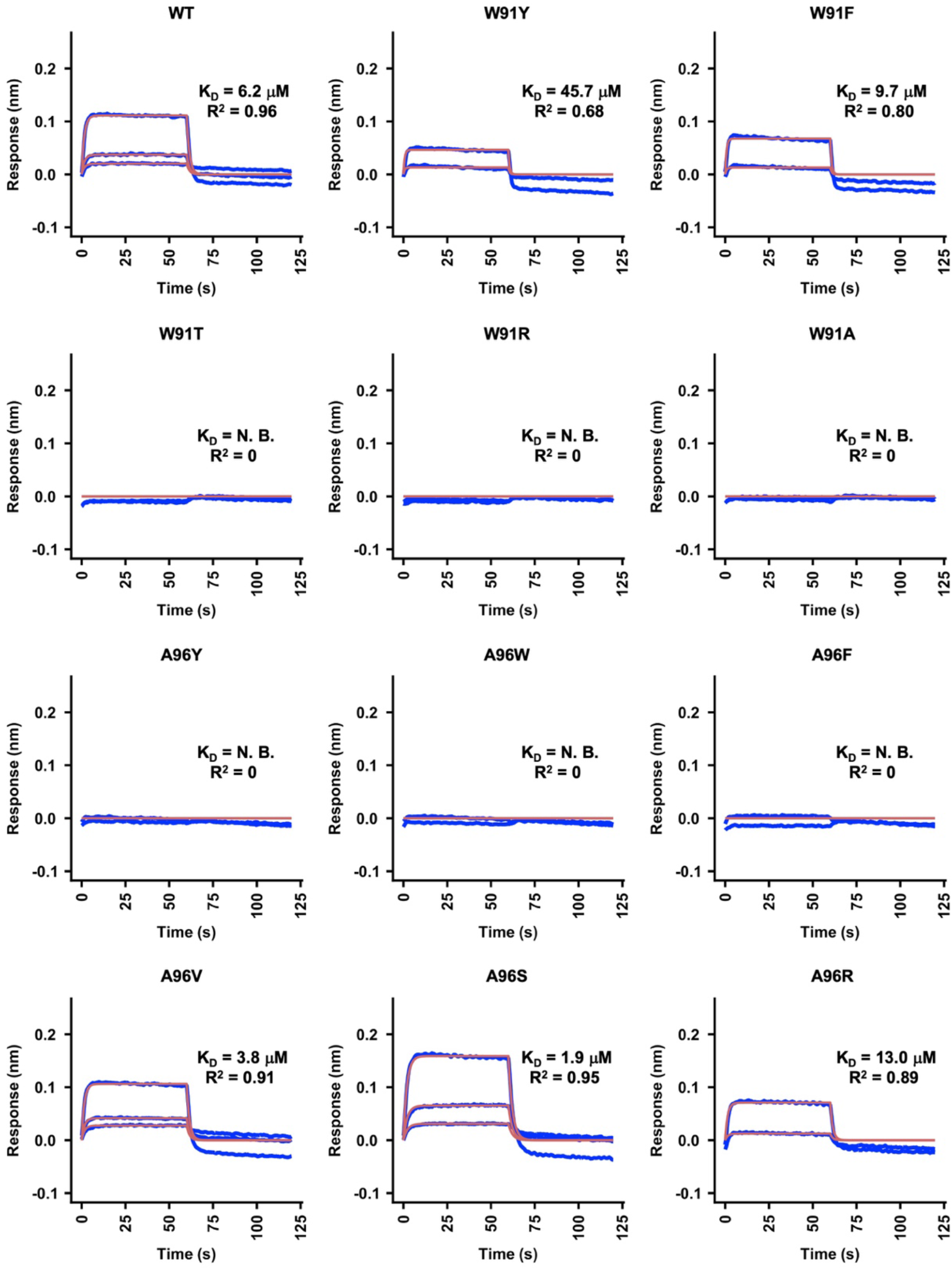
Sensorgrams for binding of CR9114 gHC mutants to mini-HA. (A) Binding kinetics of different Fabs against mini-HA were measured by biolayer interferometry (BLI). Y-axis represents the response. Blue lines represent the response curve and red lines represent a 1:1 binding model. Binding kinetics were measured for two to three Fab concentrations. N.B. indicates no binding. Dissociation constant (K_D_) and the goodness of model fitting (R^2^) are indicated.

**Supplementary Figure 3.**
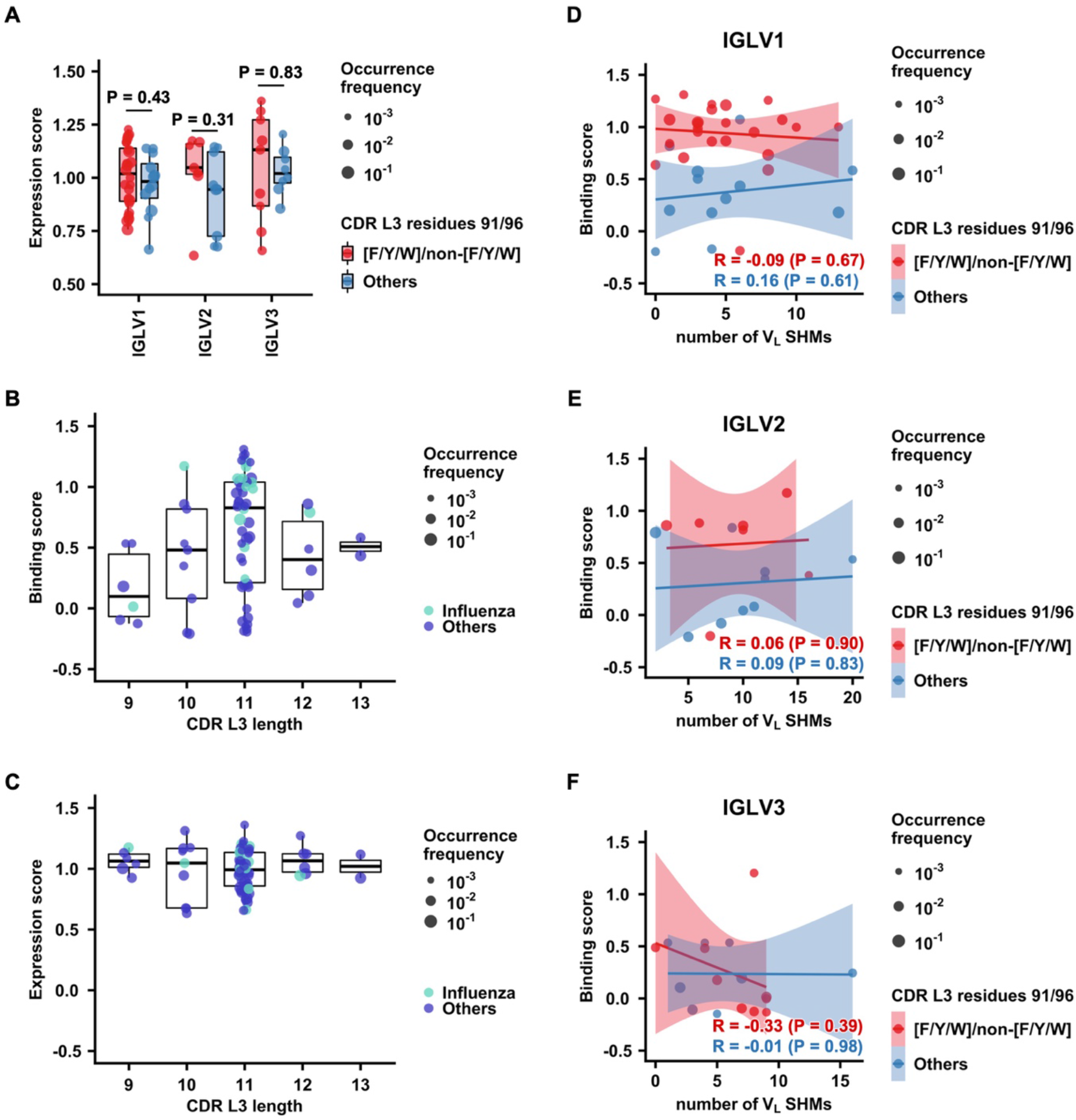
Impact of CDR L3 length and somatic hypermutation (SHM) on HA stem binding activity of CR9114. (A) Expression scores of light chain variants in different IGLV families with and without V_L_ 91_F/Y/W_/96_non-F/Y/W_ are compared. Red: with V_L_ 91_F/Y/W_/96_non-F/Y/W_; Blue: without V_L_ 91_F/Y/W_/96_non-F/Y/W_. P-values were computed by two-tailed Student’s t-test. **(B-C)** Binding **(B)** and expression **(C)** scores of light chain variants with different CDR L3 lengths are compared. Cyan: light chain variants from known IGHV1-69 antibodies to influenza virus. Blue: light chain variants from IGHV1-69 antibodies to non-influenza antigens. **(D-F)** Correlation between binding score and the number V_L_ SHM is shown for light chain variants in different light chain families, namely IGLV1 **(D)**, IGLV2 **(E)**, and IGLV3 **(F)**. The Pearson correlation coefficient (R) is indicated. P-values were computed by Pearson correlation test.

**Supplementary Figure 4.**
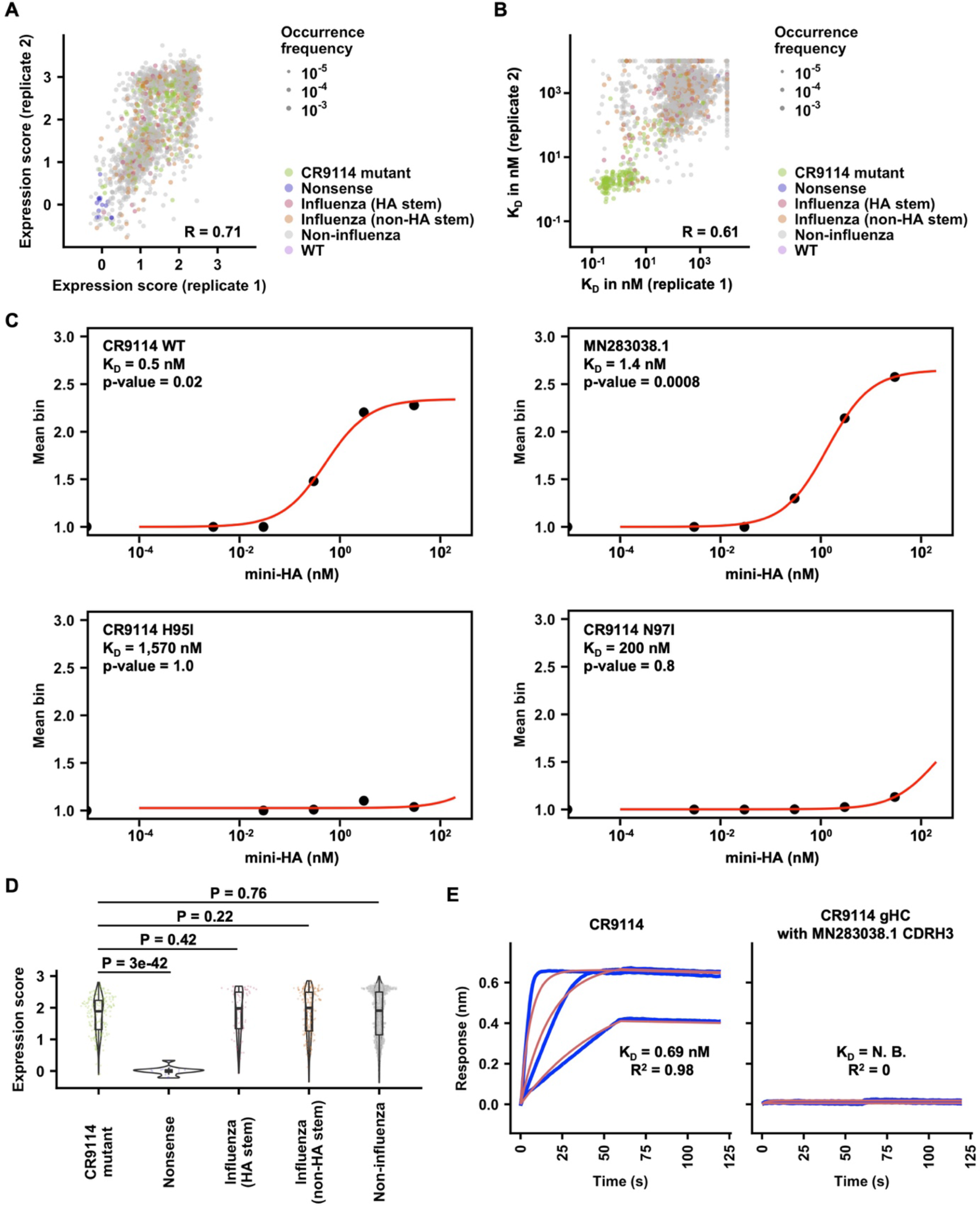
Reproducibility and analysis of Tite-Seq for CDR H3 variants. (A) Correlation of expression scores between two independent biological replicates is shown. **(B)** Correlation of apparent dissociation constant (K_D_) values between two independent biological replicates is shown. **(C)** Example titration curves inferred from Tite-Seq data. **(D)** Distributions of expression scores for CDR H3 variants from different types of antibodies. Greenish yellow: CR9114 single amino acid mutants; purple: nonsense variants with stop codons; red: CDR H3 variants from IGHV1-69 HA stem antibodies; orange: CDR H3 variants from IGHV1-69 non-HA stem influenza antibodies; grey: CDR H3 variants from IGHV1-69 non-influenza antibodies; pink: WT. **(E)** Binding kinetics of a CDR H3 variant from an IGHV1-69 non-influenza antibody (Genbank ID: MN283038.1) and CR9114 (positive control) to mini-HA were measured by BLI. Y-axis represents the response. Blue lines represent the response curve and red lines represent a 1:1 binding model. Binding kinetics were measured for at least two Fab concentrations. N.B. indicates no binding. Dissociation constant (K_D_) and the goodness of model fitting (R^2^) are indicated.

**Supplementary Figure 5.**
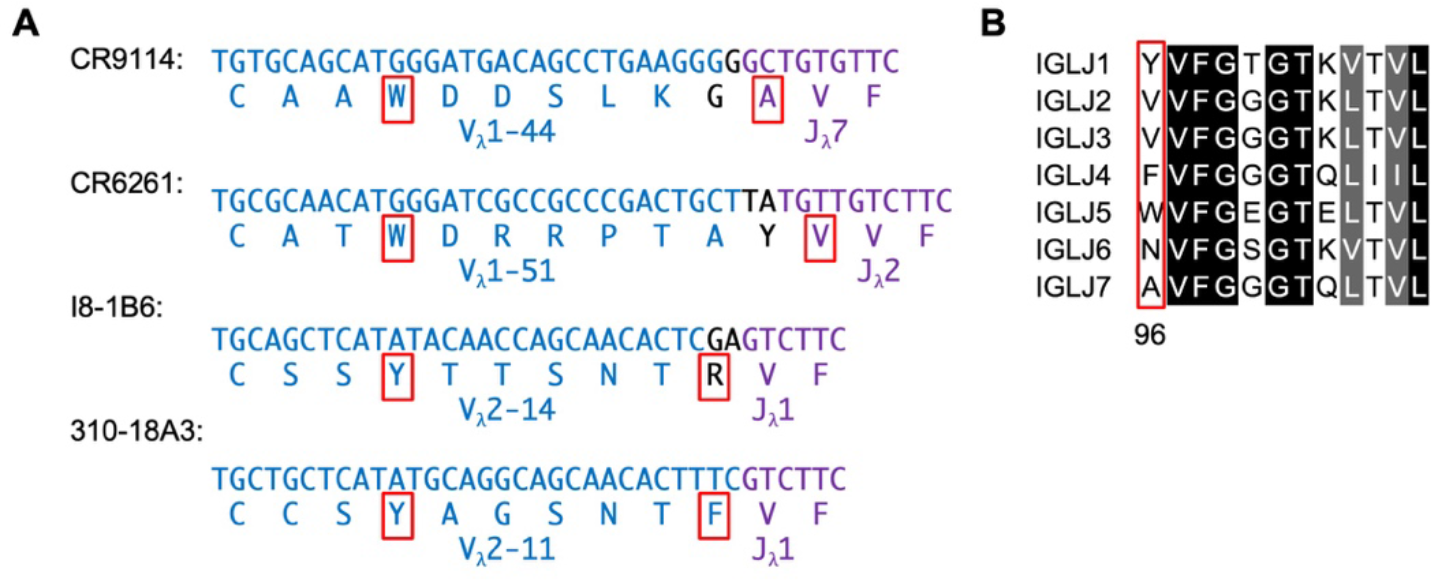
V_L_ residue 96 is sometimes encoded by IGLJ. (A) Nucleotide and amino acid sequences of light chain V-J junction are shown for different IGHV1-69 HA stem antibodies. V_L_ residues 91 and 96 are indicated in red. Blue: V-region; purple: J-region; black: N- region. **(B)** CDR L3 sequences among different IGLJ families. V_L_ residue 96 is indicated in red.

**Supplementary Table 3.**
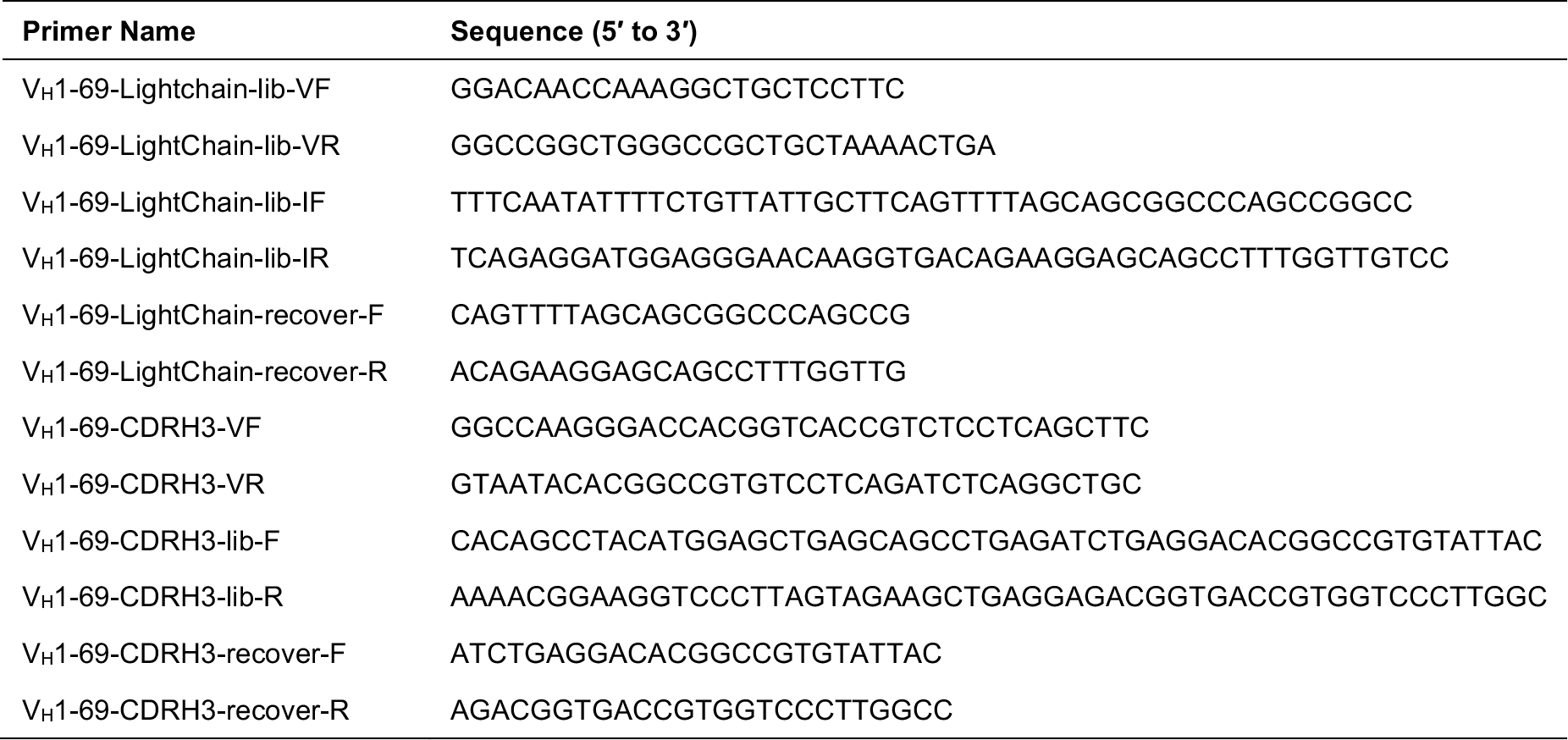
List of primers used in this study.

## REFERENCES

1. Petrova, V.N., and Russell, C.A. (2018). The evolution of seasonal influenza viruses. Nature Reviews Microbiology 16, 47–60.

2. Frentzel, E., Jump, R.L., Archbald-Pannone, L., Nace, D.A., Schweon, S.J., Gaur, S., Naqvi, F., Pandya, N., and Mercer, W. (2020). Recommendations for mandatory influenza vaccinations for health care personnel from AMDA’s Infection Advisory Subcommittee. Journal of the American Medical Directors Association 21, 25–28.e22.

3. Tricco, A.C., Chit, A., Soobiah, C., Hallett, D., Meier, G., Chen, M.H., Tashkandi, M., Bauch, C.T., and Loeb, M. (2013). Comparing influenza vaccine efficacy against mismatched and matched strains: a systematic review and meta-analysis. BMC Medicine 11, 1–19.

4. Carrat, F., and Flahault, A. (2007). Influenza vaccine: the challenge of antigenic drift. Vaccine 25, 6852–6862.

5. Impagliazzo, A., Milder, F., Kuipers, H., Wagner, M.V., Zhu, X., Hoffman, R.M., van Meersbergen, R., Huizingh, J., Wanningen, P., and Verspuij, J. (2015). A stable trimeric influenza hemagglutinin stem as a broadly protective immunogen. Science 349, 1301–1306.

6. Yassine, H.M., Boyington, J.C., McTamney, P.M., Wei, C.-J., Kanekiyo, M., Kong, W.-P., Gallagher, J.R., Wang, L., Zhang, Y., and Joyce, M.G. (2015). Hemagglutinin-stem nanoparticles generate heterosubtypic influenza protection. Nature Medicine 21, 1065–1070.

7. Dreyfus, C., Laursen, N.S., Kwaks, T., Zuijdgeest, D., Khayat, R., Ekiert, D.C., Lee, J.H., Metlagel, Z., Bujny, M.V., and Jongeneelen, M. (2012). Highly conserved protective epitopes on influenza B viruses. Science 337, 1343–1348.

8. Andrews, S.F., Graham, B.S., Mascola, J.R., and McDermott, A.B. (2018). Is it possible to develop a “universal” influenza virus vaccine? Immunogenetic considerations underlying B- cell biology in the development of a pan-subtype influenza A vaccine targeting the hemagglutinin stem. Cold Spring Harbor Perspectives in Biology 10, a029413.

9. Throsby, M., van den Brink, E., Jongeneelen, M., Poon, L.L., Alard, P., Cornelissen, L., Bakker, A., Cox, F., van Deventer, E., and Guan, Y. (2008). Heterosubtypic neutralizing monoclonal antibodies cross-protective against H5N1 and H1N1 recovered from human IgM+ memory B cells. PLOS One 3, e3942.

10. Pappas, L., Foglierini, M., Piccoli, L., Kallewaard, N.L., Turrini, F., Silacci, C., Fernandez- Rodriguez, B., Agatic, G., Giacchetto-Sasselli, I., and Pellicciotta, G. (2014). Rapid development of broadly influenza neutralizing antibodies through redundant mutations. Nature 516, 418–422.

11. Sui, J., Hwang, W.C., Perez, S., Wei, G., Aird, D., Chen, L.-m., Santelli, E., Stec, B., Cadwell, G., and Ali, M. (2009). Structural and functional bases for broad-spectrum neutralization of avian and human influenza A viruses. Nature Structural & Molecular Biology 16, 265–273.

12. Ekiert, D.C., Bhabha, G., Elsliger, M.-A., Friesen, R.H., Jongeneelen, M., Throsby, M., Goudsmit, J., and Wilson, I.A. (2009). Antibody recognition of a highly conserved influenza virus epitope. Science 324, 246–251.

13. Lang, S., Xie, J., Zhu, X., Wu, N.C., Lerner, R.A., and Wilson, I.A. (2017). Antibody 27F3 broadly targets influenza A group 1 and 2 hemagglutinins through a further variation in V_H_1- 69 antibody orientation on the HA stem. Cell Reports 20, 2935–2943.

14. Avnir, Y., Tallarico, A.S., Zhu, Q., Bennett, A.S., Connelly, G., Sheehan, J., Sui, J., Fahmy, A., Huang, C.-y., and Cadwell, G. (2014). Molecular signatures of hemagglutinin stem- directed heterosubtypic human neutralizing antibodies against influenza A viruses. PLoS Pathogens 10, e1004103.

15. Chen, F., Tzarum, N., Wilson, I.A., and Law, M. (2019). V_H_1-69 antiviral broadly neutralizing antibodies: genetics, structures, and relevance to rational vaccine design. Current Opinion in Virology 34, 149–159.

16. Xu, R., Krause, J.C., McBride, R., Paulson, J.C., Crowe Jr, J.E., and Wilson, I.A. (2013). A recurring motif for antibody recognition of the receptor-binding site of influenza hemagglutinin. Nature Structural & Molecular Biology 20, 363–370.

17. Lee, P.S., Ohshima, N., Stanfield, R.L., Yu, W., Iba, Y., Okuno, Y., Kurosawa, Y., and Wilson, I.A. (2014). Receptor mimicry by antibody F045–092 facilitates universal binding to the H3 subtype of influenza virus. Nature Communications 5, 3614.

18. Phillips, A.M., Lawrence, K.R., Moulana, A., Dupic, T., Chang, J., Johnson, M.S., Cvijovic, I., Mora, T., Walczak, A.M., and Desai, M.M. (2021). Binding affinity landscapes constrain the evolution of broadly neutralizing anti-influenza antibodies. eLife 10, e71393.

19. Boyd, S.D., Gaëta, B.A., Jackson, K.J., Fire, A.Z., Marshall, E.L., Merker, J.D., Maniar, J.M., Zhang, L.N., Sahaf, B., and Jones, C.D. (2010). Individual variation in the germline Ig gene repertoire inferred from variable region gene rearrangements. The Journal of Immunology 184, 6986–6992.

20. Lerner, R.A. (2011). Rare antibodies from combinatorial libraries suggests an SOS component of the human immunological repertoire. Molecular BioSystems 7, 1004–1012.

21. Erbelding, E.J., Post, D.J., Stemmy, E.J., Roberts, P.C., Augustine, A.D., Ferguson, S., Paules, C.I., Graham, B.S., and Fauci, A.S. (2018). A universal influenza vaccine: the strategic plan for the National Institute of Allergy and Infectious Diseases. The Journal of Infectious Diseases 218, 347–354.

22. Nachbagauer, R., Feser, J., Naficy, A., Bernstein, D.I., Guptill, J., Walter, E.B., Berlanda- Scorza, F., Stadlbauer, D., Wilson, P.C., and Aydillo, T. (2021). A chimeric hemagglutinin- based universal influenza virus vaccine approach induces broad and long-lasting immunity in a randomized, placebo-controlled phase I trial. Nature Medicine 27, 106–114.

23. Andrews, S.F., Cominsky, L.Y., Shimberg, G.D., Gillespie, R.A., Gorman, J., Raab, J.E., Brand, J., Creanga, A., Gajjala, S.R., and Narpala, S. (2023). An influenza H1 hemagglutinin stem-only immunogen elicits a broadly cross-reactive B cell response in humans. Science Translational Medicine 15, eade4976.

24. Adams, R.M., Mora, T., Walczak, A.M., and Kinney, J.B. (2016). Measuring the sequence- affinity landscape of antibodies with massively parallel titration curves. eLife 5, e23156.

25. Starr, T.N., Greaney, A.J., Hilton, S.K., Ellis, D., Crawford, K.H., Dingens, A.S., Navarro, M.J., Bowen, J.E., Tortorici, M.A., and Walls, A.C. (2020). Deep mutational scanning of SARS-CoV-2 receptor binding domain reveals constraints on folding and ACE2 binding. Cell 182, 1295–1310.e1220.

26. Chao, G., Lau, W.L., Hackel, B.J., Sazinsky, S.L., Lippow, S.M., and Wittrup, K.D. (2006). Isolating and engineering human antibodies using yeast surface display. Nature Protocols 1, 755–768.

27. Benson, D.A., Cavanaugh, M., Clark, K., Karsch-Mizrachi, I., Lipman, D.J., Ostell, J., and Sayers, E.W. (2012). GenBank. Nucleic Acids Research 41, D36–D42.

28. Ye, J., Ma, N., Madden, T.L., and Ostell, J.M. (2013). IgBLAST: an immunoglobulin variable domain sequence analysis tool. Nucleic Acids Research 41, W34–W40.

29. Benatuil, L., Perez, J.M., Belk, J., and Hsieh, C.-M. (2010). An improved yeast transformation method for the generation of very large human antibody libraries. Protein Engineering, Design and Selection 23, 155–159.

30. Ekiert, D.C., Friesen, R.H., Bhabha, G., Kwaks, T., Jongeneelen, M., Yu, W., Ophorst, C., Cox, F., Korse, H.J., and Brandenburg, B. (2011). A highly conserved neutralizing epitope on group 2 influenza A viruses. Science 333, 843–850.

31. Mölder, F., Jablonski, K.P., Letcher, B., Hall, M.B., Tomkins-Tinch, C.H., Sochat, V., Forster, J., Lee, S., Twardziok, S.O., and Kanitz, A. (2021). Sustainable data analysis with Snakemake. F1000Research *10*.

32. Zhang, J., Kobert, K., Flouri, T., and Stamatakis, A. (2014). PEAR: a fast and accurate Illumina Paired-End reAd mergeR. Bioinformatics 30, 614–620.

33. Starr, T.N., Greaney, A.J., Hilton, S.K., Ellis, D., Crawford, K.H., Dingens, A.S., Navarro, M.J., Bowen, J.E., Tortorici, M.A., and Walls, A.C. (2020). Deep mutational scanning of SARS-CoV-2 receptor binding domain reveals constraints on folding and ACE2 binding. Cell 182, 1295–1310. e1220.

34. Tareen, A., and Kinney, J.B. (2020). Logomaker: beautiful sequence logos in Python. Bioinformatics 36, 2272–2274.

35. Wu, N.C., Grande, G., Turner, H.L., Ward, A.B., Xie, J., Lerner, R.A., and Wilson, I.A. (2017). In vitro evolution of an influenza broadly neutralizing antibody is modulated by hemagglutinin receptor specificity. Nature Communications 8, 15371.

